# Parthanatos-inducing zinc agent C010DS-Zn elicits anti-tumor immune responses involving T cells and macrophages *in vivo*

**DOI:** 10.1101/2021.03.18.433812

**Authors:** Jinhyuk Fred Chung, Zhisheng Her, Wai Mun Kong, Qingfeng Chen

**Affiliations:** Xylonix PTE. LTD., Singapore 079906; Invivocue PTE. LTD., Singapore; Humanized Mouse Unit, IMCB, A*Star, Singapore; Invitrocue PTE. LTD., Singapore

## Abstract

Immune checkpoint inhibitors opened a new horizon in cancer therapy by enabling durable and complete responses in patients, but their wider application against general solid cancers has been hampered by the lack of a broadly acting anti-cancer immune response initiating agents. Parthanatos is a previously unexplored immunogenic programmed necrosis mechanism that is triggered by the hyperactivation of PARP DNA repair and executed by an efficient DNA-fragmentation mechanism. We developed a proprietary macromolecular zinc complex agent C010DS-Zn that efficiently induced parthanatos against 4T1 murine cancer cells *in vitro*, which was characterized as PARP-mediated necrotic death with massive DNA damages. *Ex vivo* screening of its cytotoxicity against a panel of 53 low-passage human solid cancer PDX tumor fragments demonstrated its consistent delivery of characteristically DNA-damaging cell death that was unseen in the corresponding apoptosis positive controls. Further characterization of its *in vivo* treatment effects versus the immunosuppressive 4T1-Balb/c and immunogenic CT26-Balb/c syngeneic cancer models showed that sufficiently high intravenous C010DS-Zn treatments led to robust initiation of the tumor-suppressed antitumor immune compartments such as T cells and macrophages. At lower non-anticancer doses, C010DS-Zn treatment still led to significantly reduced macrophage content and inflammation in the 4T1 tumor, suggesting its potential utility against macrophage-mediated inflammations such as those seen in MIS-C or COVID19. Given the observation of its low serum bioavailability in a rat pharmacokinetic study, these results suggest potential development opportunities for C010DS-Zn to become a widely applicable immune initiation agent with chemo-like broad applicability upon its pharmacokinetic improvements.

## Introduction

Adding a neo-antigen targeting cancer vaccination for initiating anti-cancer T cell immune response is a mainstay strategy being investigated for improving the effectiveness of current immunotherapies versus a wider range of cancer indications. While there have been successes in this investigational approach across several vaccine platforms (*1*), a challenging drawback of this approach is the narrow target coverage of each vaccine candidate due to high neo-antigen heterogeneity within and across cancer types that escalate the number and the cost of vaccine developments needed for treating common cancers. As the increasing cost of combination cancer immunotherapy is becoming a major hurdle against its wider use (*2*), the need for a broadly applicable anti-cancer immune response initiator is emerging.

Cancer-targeted induction of immunogenic cell death such as necroptosis (*3*), pyroptosis (*4*), or parthanatos (*5*) is an alternative immune response initiation strategy wherein the killed cancer cells become the immunogenic template for their own neo-antigen presentation. Its potential has been demonstrated in the leading preclinical studies of necroptosis, wherein the vaccination using *in vitro* RIPK3-mRNA necroptized cancer cells led to robust dendritic cell (DC) maturation and CD8+ T cell cross priming with remarkable antitumor effects *in vivo* (*3*). The same group further demonstrated that *in vivo* necroptosis of xenografted tumor using electroporation-delivered MLKL-mRNA (*6*) conferred immune initiation and anti-tumor benefits, highlighting its potential for pharmaceutical translation. Similar anti-cancer immunity effects from the use of pyroptosis or parthanatos, on the other hand, have not been reported yet.

Parthanatos is a mode of programmed necrosis triggered by the DNA damage sensor/repair enzyme PARP hyperactivation (*7*), wherein the resulting accumulation of the reaction product PAR-polymer leads to downstream death execution events of nuclear AIF translocation and subsequent DNA fragmentation (*8*). Acute toxicities of zinc (*9*) or cadmium (*10*) were previously reported to involve PARP1 activity, hinting possibilities of their use in parthanatos induction against cancer. We have developed a degradable ligand-modified biopolymer-based zinc (II) complex agent C010DS-Zn (a Cy5.5 labelled version of 010DS-Zn for investigative use) for selective induction of parthanatos targeting cancer. Here we report its *in vitro* mode-of-action study results using 4T1 murine triple negative breast cancer cell line in corroborating its efficient parthanatos-induction activity, its *ex-vivo* screening results against 53 patient-derived solid cancer models in assessing its broad application potential, and *in vivo* test results against the immunogenic murine cancer CT26 and the immunosuppressive murine cancer 4T1 in describing its respective immunotherapeutic activities.

## Results

### C010DS-Zn induced more potent necrosis versus control compounds in vitro

The studied C010DS-Zn was zinc-carrying gamma-polyglutamate conjugated with 1 Cy5.5 fluorescent reporter, 2 folate-(ester)-PEG4-NH3, 2 cRGDfK-(ester)-PEG4-NH3, and 10 pyrithione-(disulfide)-PEG3-NH3 per 32 kDa (Mw) polymer backbone, with the zinc loaded at 7 Zn^2+^ to 1 pyrithione (Pyr) sidechain molar ratio via backbone complexation (*11*) (**Figure 1B**). *In vitro* cytotoxicity of C010DS-Zn and relevant control groups including zinc sulfate, zinc sulfate-sodium Pyr mixture at 7 Zn^2+^ to 1 Pyr molar ratio modelling the C010DS-Zn payload (7Zn-1Pyr), and C005D-Zn (a positive control compound with identical structure and zinc content minus the Pyr-sidechain modifications)(**Figure 1B**) were therefore tested.

**Figure 1.**
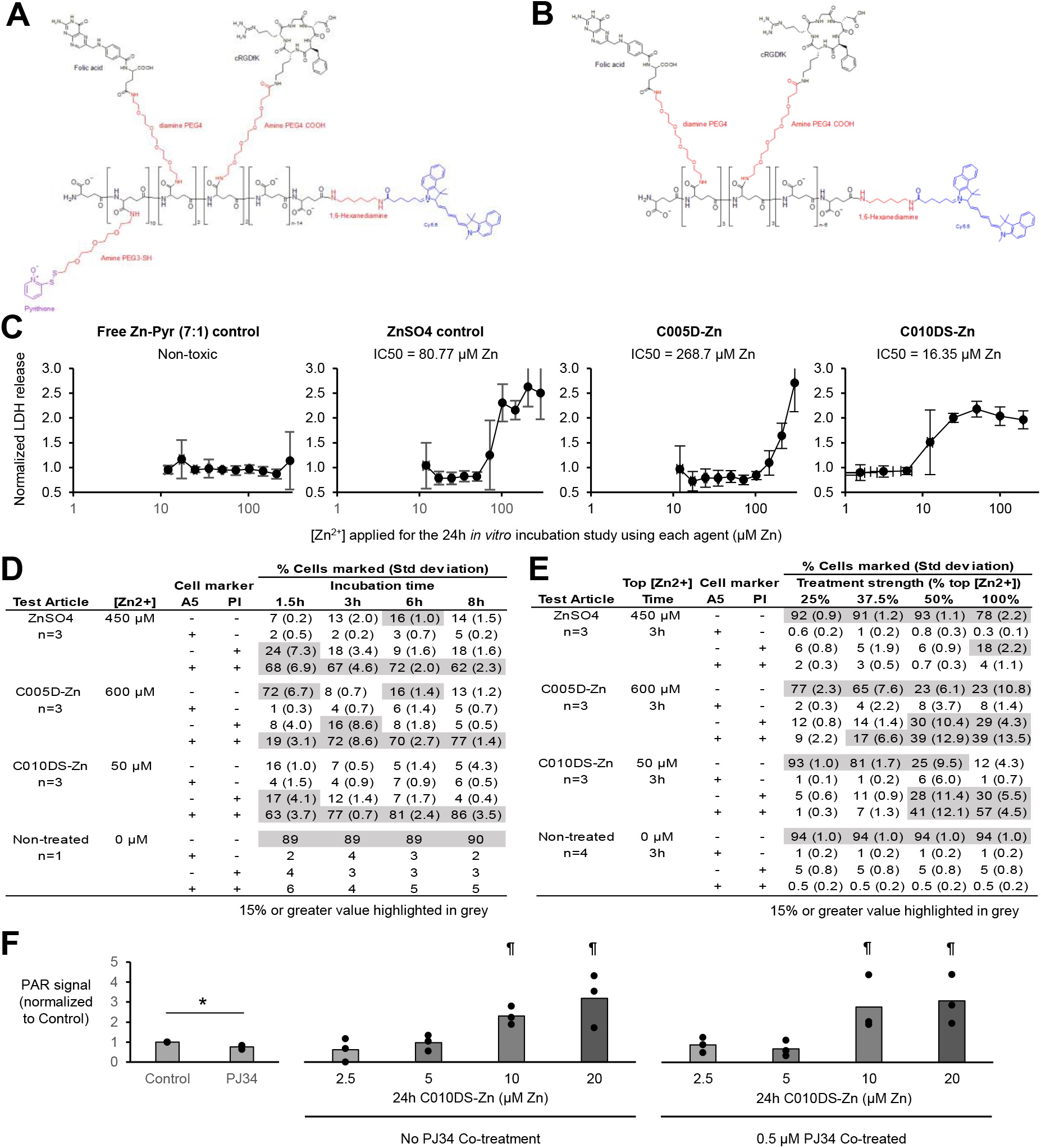
Basic substance information on C010DS-Zn, and its control substances C005D-Zn, zinc sulfate (ZnSO_4_), and “Free Zn-Pyr (7:1)” (the mixture of zinc sulfate and sodium pyrithione at zinc-to-pyrithione molar ratio of 7-to-1). **(A)** Molecular structure of C010DS, 34 kDa (M_w_) γPGA polymer each conjugated with 1 Cy5.5 label, 2 PEG-folate sidechains, 2 PEG-cRGDfK sidechains, and 10 PEG-S-S-pyrithione sidechains. C010DS-Zn preparation is described in the Methods. **(B)** Molecular structure of C005D, 34 kDa (M_w_) γPGA polymer each conjugated with 1 Cy5.5 label, 3 PEG-folate sidechains, and 3 PEG-cRGDfK sidechains. C005D-Zn preparation is described in the Methods. **(C)** Comparative *in vitro* cytotoxicity evaluation on the tested compounds using LDH release assays after 24h treatments against 4T1 cells. **(D)** Results of the *in vitro* time-resolved apoptosis-necrosis flow cytometry assay at fixed doses against 4T1 cells using A5 and PI labels. **(E)** Results of the *in vitro* dose-resolved apoptosis-necrosis flow cytometry assay at 3h time point against 4T1 cells using A5 and PI labels. **(F)** *In vitro* PAR-ELISA assay results on the C010DS-Zn treated 4T1 cells with or without the PARP inhibitor PJ34. \**p*<0.05. ^¶^Indicated group displayed significantly higher PAR signal than the Control, 2.5 μM Zn, or 5 μM Zn C010DS-Zn treatment groups at *p*<0.05 with or without PJ34 co-treatment.

24h *in vitro* cytotoxicity was tested for each compound against 4T1 murine triple negative breast cancer cell line in obtaining their IC50_24h_ values using lactate dehydrogenase release assay (LDH assay), which were expressed in zinc concentration (μM Zn) units for comparative viewing. C010DS-Zn showed markedly higher potency at IC50_24h_ of 16.35 μM Zn versus zinc sulfate (80.7 μM Zn) or C005D-Zn (268.7 μM Zn). 7Zn-1Pyr did not show cytotoxic response in the concentration range tested (12.1 μM Zn – 300 μM Zn) (**Figure 1C**).

Flow cytometry analyses using annexinV (A5) and propidium iodide (PI) markers were performed to compare the *in vitro* apoptotic/necrotic cell death feature of the observed cytotoxic effects around IC50_24h_-defined concentration gradient after 3h treatment (**Figure 1E**), or over time at an excess concentration over IC50_24h_ (**Figure 1D**). 7Zn-1Pyr was excluded from the test as its IC50_24h_ value could not be derived due to its lack of cytotoxicity. The time resolved study treatment showed that C010DS-Zn (50 μM Zn) and zinc sulfate (450 μM Zn) groups induced mainly PI+A5+ and minor PI+ only necrotic cell death 1.5h after the treatment, while C005D-Zn (600 μM Zn) showed similar necrotic death 3h after the treatment. A5+ only apoptotic death response was not observed in any of the test groups or at any time points. The concentration resolved study after 3h treatment showed that C010DS-Zn, C005D-Zn, and zinc sulfate respectively induced up to 77%, 62%, 26% PI+ only or PI+A5+ necrotic cell death in concentration-dependent manners without notable increases in A5+ only apoptotic cell deaths. Also, the proportion of PI+A5+ versus PI+ only necrosis was found highest in the C010DS-Zn (57% vs 30%) group and the lowest in the zinc sulfate group (4% vs 18%). Collectively, these cytotoxicity tests demonstrated C010DS-Zn as the most potent and efficient inducer of the PI+A5+ necrotic cell death per given zinc content against the 4T1 cells.

### In vitro mode-of-action studies using 4T1 murine cancer cell line corroborate rapid and potent parthanatos induction by C010DS-Zn

In addition to the cytotoxicity featuring the simultaneous uptake of PI and A5, Fatokun et al. previously defined PARP-dependent nuclear translocation of AIF and subsequent generation of massive DNA breaks as the hallmark features of parthanatos (*8*). To investigate whether the observed PI+A5+ necrotic 4T1 cell deaths by the zinc compounds (**Figure 1D and 1E**) accompanied these features, we performed confocal microscopy studies after 1.5h treatments for assessing the occurrence of PARP-dependent cytotoxicity, nuclear AIF translocation, and DNA breaks using fluorescent probes of anti-AIF, TUNEL, and Hoechst 33342 (Hoechst) DNA stain upon co-incubation with a potent PARP1 and PARP2 inhibitor PJ34. Assessment of PAR-polymer content, however, could not be incorporated into this experiment due to unavailability of sufficiently sensitive fluorescent probe for PAR-polymer during this study. Also, any longer treatment time could not be applied to this experiment due to extensive cytotoxic changes observed in the C010DS-Zn treated groups.

After the 1.5h treatment, zinc sulfate and C005D-Zn treatment did not cause marked cell detachment across all concentrations tested, although the cell morphologies in the higher concentration treatments showed cytotoxic changes. C010DS-Zn treatment, however, led to complete cell loss in the 60 μM Zn treated group and extensive cell detachments even in the 15 μM Zn treated group. Co-treatment with 0.5 μM PJ34 (a selective PARP1 and PARP2 inhibitor) significantly attenuated the cell detachments in the C010DS-Zn 15 μM Zn group but showed no protective effects in the 60 μM Zn treated group (**Figure 2B**).

**Figure 2.**
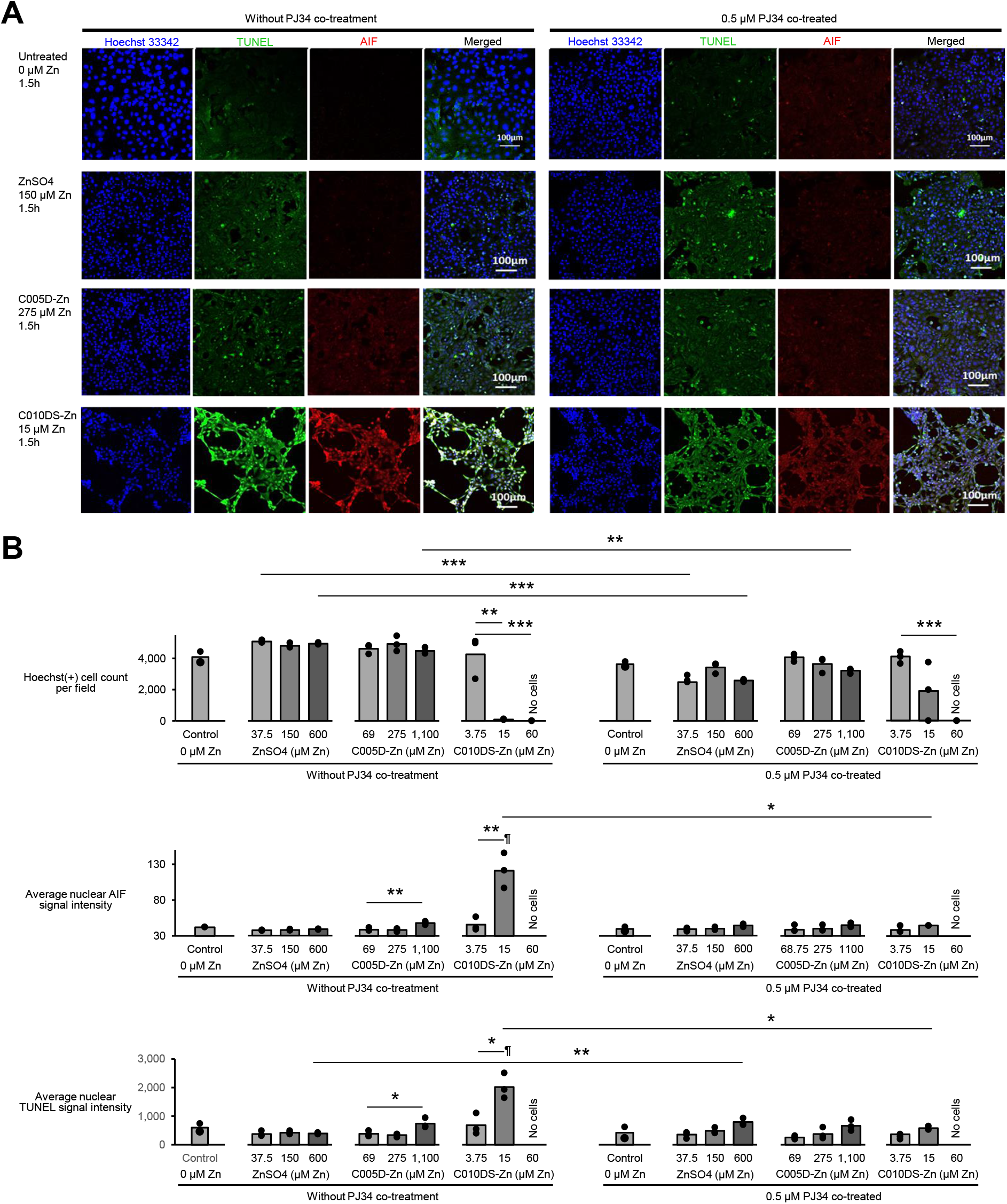
*In vitro* attenuation of 1.5h C010DS-Zn treatment effects against 4T1 including nuclear AIF translocation, DNA fragmentation, and cell viability loss by PARP inhibitor PJ34. **(A)** Representative confocal fluorescence images of the 4T1 cells treated for 1.5h with vehicle, ZnSO4, C005D-Zn, or C010DS-Zn at different concentrations, labelled with Hoechst (blue: adherent cell count), TUNEL (green: DNA breaks), and anti-AIF (red), with or without the PARP inhibitor PJ34 co-incubation. Bar=100μm. **(B)** Quantitative analysis of the fluorescence images for cell viability, nuclear AIF translocation, and nuclear TUNEL intensity using CirAvgInten function. 3 imaging fields quantified for each group (n=3). \**p*<0.05. \*\**p*<0.005. \*\*\**p*<0.0005. ¶Indicated group’s average value is significantly greater than those of all other treatment groups minimally at *p*<0.05. C010DS-Zn at 60 μM Zn treatment group resulted in complete cell loss, and hence its quantitative imaging analyses could not be performed.

Subsequent quantitative imaging analysis on the averaged nuclear AIF signal and TUNEL signals, respectively representing the extent of nuclear AIF translocation and the resulting DNA damages in the detected nuclei, showed that C010DS-Zn caused significant nuclear AIF translocation and massive DNA damage induction at 15 μM Zn, which was abrogated by the PJ34 co-treatment. Similar but significantly less trend was seen in the C005D-Zn treated groups. Zinc sulfate treatment, on the other hand, failed to elicit significantly higher nuclear AIF translocation nor the DNA damages.

In further validation of the PARP enzyme involvement observed in the cytotoxicity of C010DS-Zn, we performed PAR-polymer quantification using a commercial ELISA kit and observed that 24h C010DS-Zn treatment indeed produced dose-dependent accumulation of PAR-polymer indicative of PARP enzyme hyperactivity in the *in vitro* treated 4T1 cells (**Figure 1F**).

As these observations are consistent with the cell death characteristics of parthanatos as stipulated by Fatokun et al. (*8*), C010DS-Zn was collectively demonstrated as a remarkably stronger inducer of parthanatos over zinc sulfate, C005D-Zn or other control substances tested.

### Imaging-based ex vivo screening of C010DS-Zn cytotoxicity against 53 patient-derived solid

To determine the applicability and consistency of C010DS-Zn pharmacologic activity against wider solid cancer types, we employed the imaging-based *ex vivo* cytotoxicity screening service platform using the tumor fragments of patient-derived xenografts (PDX) from Champions Oncology/Phenovista. 53 PDX models from 8 cancer types including triple negative breast cancer (BrC-TNBC), non-triple negative breast cancer (BrC-nonTNBC), colorectal cancer (CRC), non-small cell lung cancer (NSCLC), ovarian cancer (OVC), pancreatic cancer (Panc), prostate cancer (PrC), and sarcoma were chosen. C010DS-Zn demonstrated cytotoxicity against all 53 models tested with IC50 values ranging from 0.07 μg Zn/mL to 1.91 μg Zn/mL, with ovarian cancers showing statistically non-significant but higher average IC50 values than other cancer types **(Figure 3A)**. Statistically significant correlation between C010DS-Zn IC50 values and each model’s tumor mutation burden (TMB) or microsatellite instability (MSI) score was also not found, indicating that C010DS-Zn cytotoxicity did not depend on the innate mutational burden of the target models (**Figure 3B**) despite the conventional view of DNA-damage mediated PARP activation as the first key step in parthanatos induction (*8*). Comparative assessment of the γH2AX level, an indicator double stranded DNA (dsDNA) breaks, in the tumor fragments between the apoptosis-inducing 10%DMSO (*12*) positive control treatment and the top-dose C010DS-Zn treatment group showed that C010DS-Zn induced markedly greater dsDNA breaks than the apoptosis positive control, even in the model with the highest IC50 value tested (**Figure 3C**). This characteristically massive induction of dsDNA breaks by C010DS-Zn treatment versus the 10%DMSO positive control treatment was further observed in 50 of the 53 ex vivo PDX models tested, indicating high consistency in its pharmacologic action versus diverse solid cancer types (**Figure 4**).

**Figure 3.**
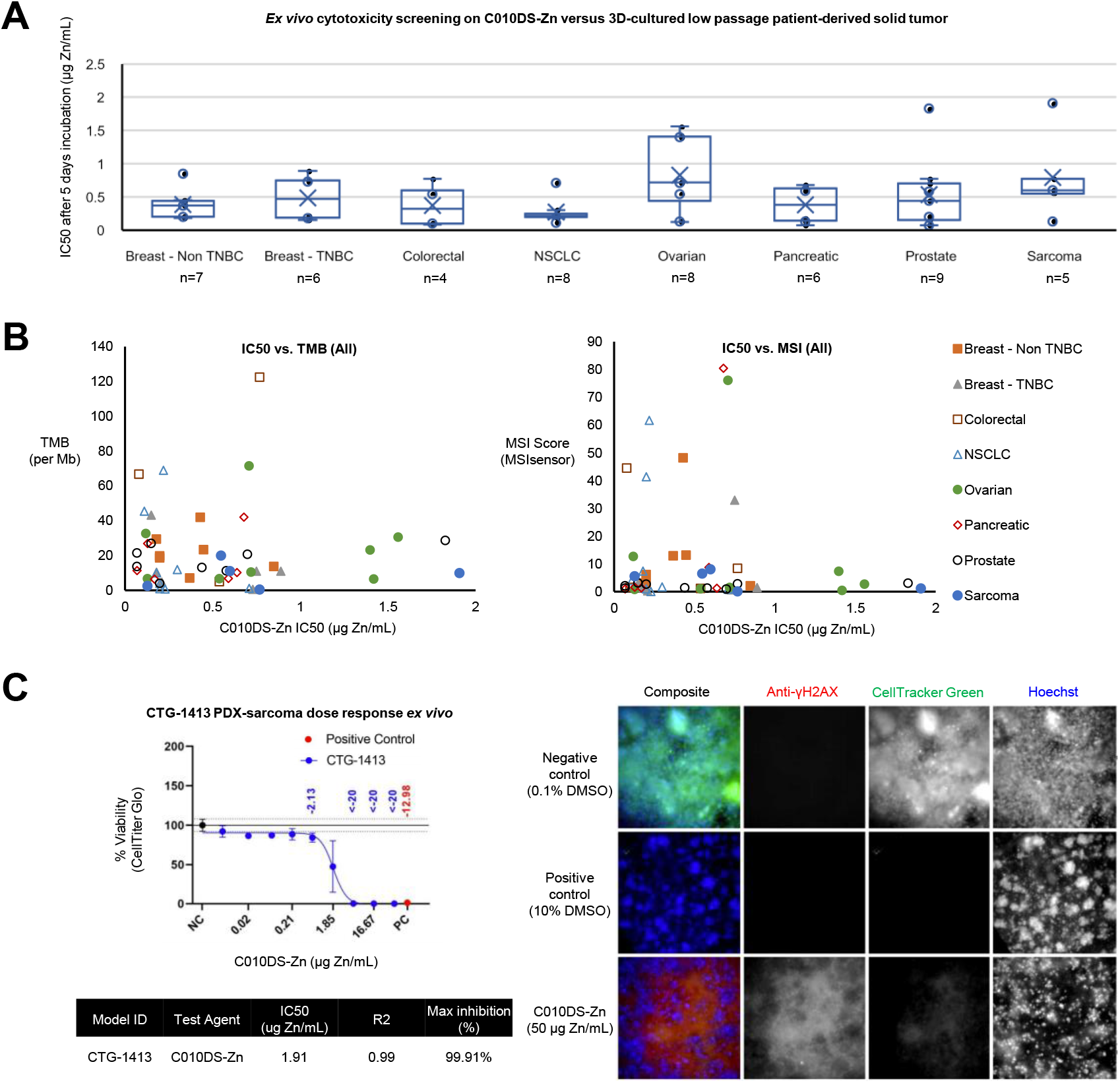
*Ex vivo* C010DS-Zn cytotoxicity screening against 53 patient-derived xenograft (PDX) tumor fragments demonstrated consistent cytotoxic action regardless of their tumor origin, TMB or MSI status. **(A)** *Ex vivo* IC50 values of C010DS-Zn versus the 53 PDX-tumor types stratified by the tumor sites and visualized using Whisker plot (minimum, lower quartile, median, x=average, upper quartile, maximum, with outliers in circles). **(B)** Plots of the *ex vivo* IC50 values versus the TMB or the MSI scores associated with each PDX-tumor fragment. **(C)** An example *ex vivo* cytotoxicity response data from the CTG-1413 sarcoma PDX-tumor fragment that showed the highest IC50 value within the panel at 1.91μg Zn/mL, together with its imaging analysis showing greater uptake of anti-γH2AX than its positive control indicating characteristically massive dsDNA breaks induced by C010DS-Zn.

**Figure 4.**
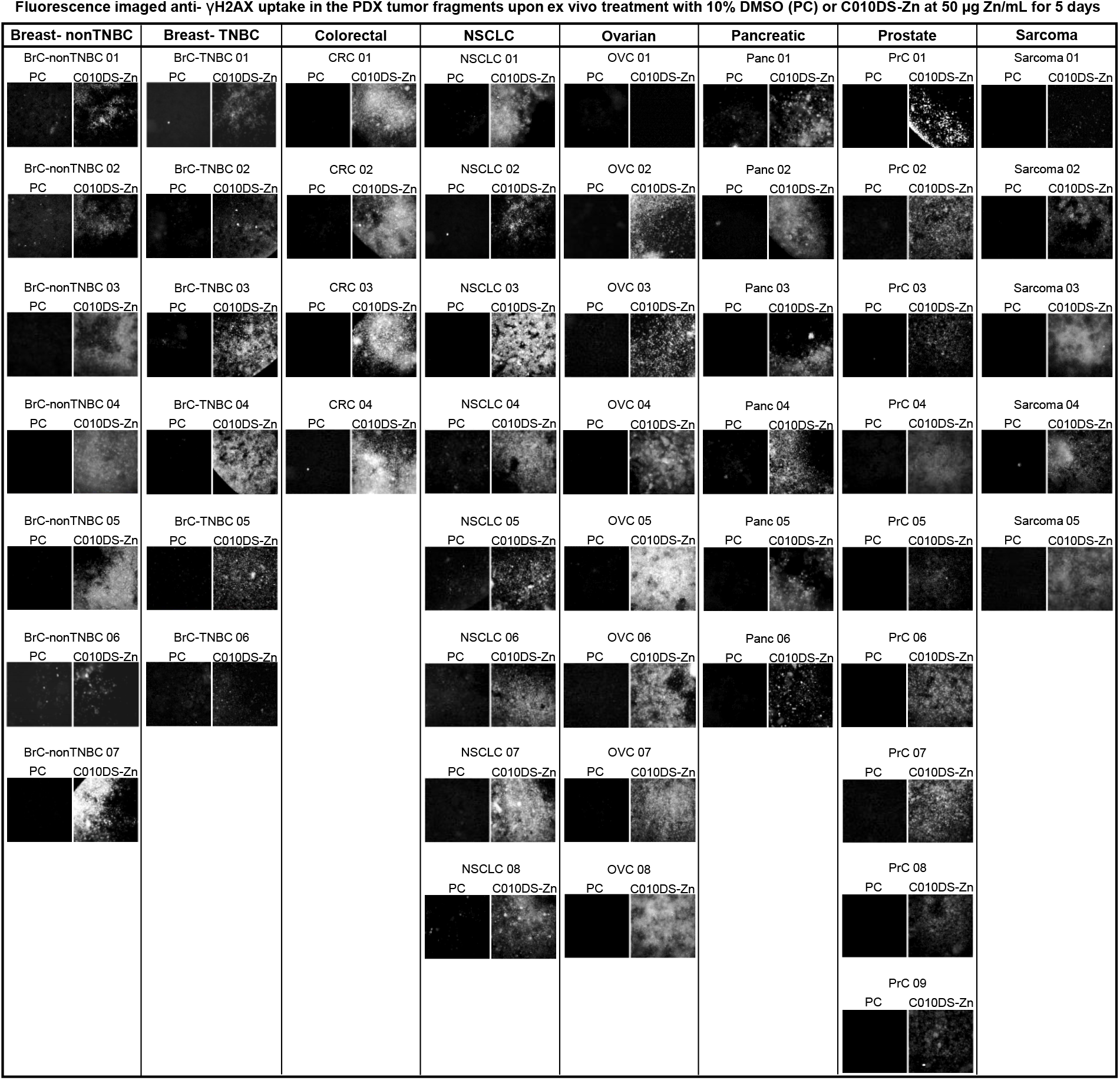
Comparative fluorescence imaging characterization of anti-γH2AX uptake in the PDX tumor fragments upon *ex vivo* treatment with 10% DMSO (PC) or C010DS-Zn at 50 μg Zn/mL for 5 days against the 53 PDX tumor types tested. Grossly increased anti-γH2AX uptake was observed in the C010DS-Zn treated groups versus the PC treated groups in 50 of the 53 PDX tumor types tested, indicating consistently greater dsDNA breaks caused by the C010DS-Zn treatment versus that by the PC treatment.

### Murine cancer models 4T1-Balb/c and CT26-Balb/c showed therapeutic response to C010DS-Zn treatments with favourable anti-cancer immune initiation profile

Next we decided to test the *in vivo* therapeutic effects of intravenously (i.v.) administered C010DS-Zn against the immunogenic syngeneic cancer model CT26-Balb/c and immunosuppressive syngeneic cancer model 4T1-Balb/c. Briefly, CT26 is a widely used immunogenic murine colorectal cancer model and a responder to anti-PD1 treatment. 4T1, on the other hand, is a widely used immunosuppressive triple negative breast cancer model that does not respond to anti-PD1 treatment (*13*).

After initial trial-and-error attempts targeting the 4T1-Balb/c model, we discovered that twice daily i.v. administration of C010DS-Zn at each injection dose of 2 mg Zn/Kg was needed in producing observable therapeutic efficacy. 50%+ tumor surface ulcerations problems were frequently encountered during pilot studies in non-treated animals, but toxicity signs meeting the General Toxicity definitions were not observed during the final study (**Figure 5A** and **Materials and Methods**). While direct comparison of the tumor growth rate was difficult due to rapid tumor ulceration developments in the vehicle control group (30 mM HEPES at pH 7), end point tumor microenvironment (TME) immune cell analysis revealed that the twice daily injection of 2mg Zn/Kg C010DS-Zn led to significant immune compartment expansions in CD4 T cells, CD 8 T cells, Granzyme B+ dendritic cells (GB+ DC), and macrophages. Particularly in the macrophage compartment, proportion of anti-tumor M1-like macrophages was significantly escalated (25% vs Control group’s 7%, *p*<0.05) while that of the pro-tumor M2-like macrophages was suppressed (18% vs Control group’s 35%, *p*<0.005) (**Figure 5A**). C010DS-Zn dosing at lesser amount or frequency including the once daily i.v. at 1mg Zn/Kg injection did not produce such results. C010DS-Zn treatments were well tolerated, and the animal body weights remained stable.

**Figure 5.**
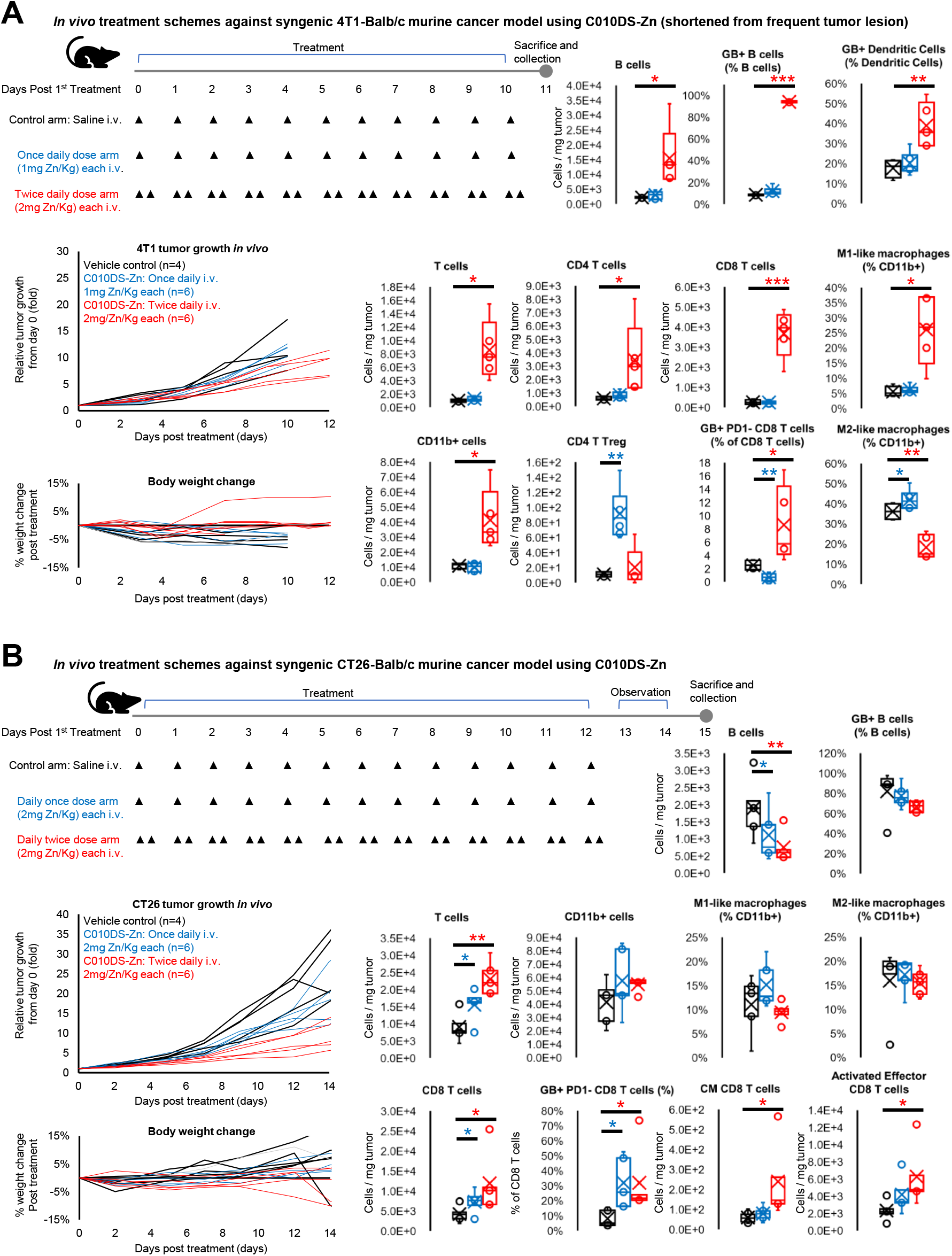
*In vivo* characterization of the anti-tumor responses induced by C010DS-Zn intravenous treatments against 4T1 murine TNBC model on Balb/c mice and against CT26 murine CRC model on Balb/c mice. **(A)** 4T1-Balb/c treatment model scheme, tumor growth kinetics, and notable immune responses in the TME. **(B)** CT26-Balb/c treatment model scheme, tumor growth kinetics, and notable immune responses in the TME. \**p*<0.05. \*\**p*<0.005. \*\*\**p*<0.0005.

C010DS-Zn i.v. treatment against CT26-Balb/c model produced better pronounced and dose-dependent responses in both tumor growth and in the end point TME immune cell analysis. No toxicity signs meeting the General Toxicity definitions were observed during the study (**Figure 5B** and **Materials and Methods**). While CT26-Balb/c model showed tumor growth reduction in both single and twice daily injection regimens unlike the 4T1-Balb/c model, significant immune response initiation effect was limited to CD 8 T cell compartment (**Figure 5B**). C010DS-Zn treatments were well tolerated, and the animal body weights remained stable.

Interestingly, when comparing the end point tumor immune profile between those of 4T1-Balb/c and CT26-Balb/c control models, we observed that the T cell (1E+3 in 4T1 vs. 9E+3 in CT26), CD11b+ myeloid cells (1E+4 in 4T1 vs. 4E+4 in CT26), and M1 macrophages levels (5% of CD11b+ cells in 4T1 vs. 11% of CD11b+ cells in CT26) were markedly suppressed in the 4T1 model versus the CT26 model. Same comparison between the twice daily C010DS-Zn treated groups of 4T1 and CT26, on the other hand, showed narrower gap between the two tumor model immune compositions in the T cells (8E+3 in 4T1 vs. 2E+4 in CT26) and CD11b+ cells (4E+4 in 4T1 vs. 5E+4 in CT26). Post treatment M2-like macrophage levels in the 4T1 and the CT26 models were similar at 18% and 15%. Collectively, these observations indicated that the anti-tumor immune initiation effects of the twice daily C010DS-Zn treatments were more pronounced in the tumor-suppressed immune compartments of 4T1-Balb/c and CT26-Balb/c models.

### Rat plasma pharmacokinetic (PK) profile of C010DS-Zn and the macrophage clearing effects of C010DS-Zn at lower doses in the TME of 4T1-Balb/c model

Our observation that twice daily i.v. treatment using C010DS-Zn was needed in producing the anti-tumor immune response initiation in 4T1-Balb/c model prompted us to test the pharmacokinetic profile of i.v. injected C010DS-Zn (**Figure 6A**). Evaluation of in-rat PK profile after a single bolus injected i.v. C010DS-Zn in non-tumor bearing Sprague-Dawley Rats revealed that only a fraction of the zinc injected by C010DS-Zn was found in the cell-free plasma compartment of blood with T_max_ of 2h, indicating rapid systemic clearance of the injected C010DS-Zn by mononuclear phagocytic system (MPS) (*14*). As MPS mainly consists of hepatic and splenic macrophages, this observation suggested possible direct cytotoxic interaction of C010DS-Zn against macrophages.

**Figure 6.**
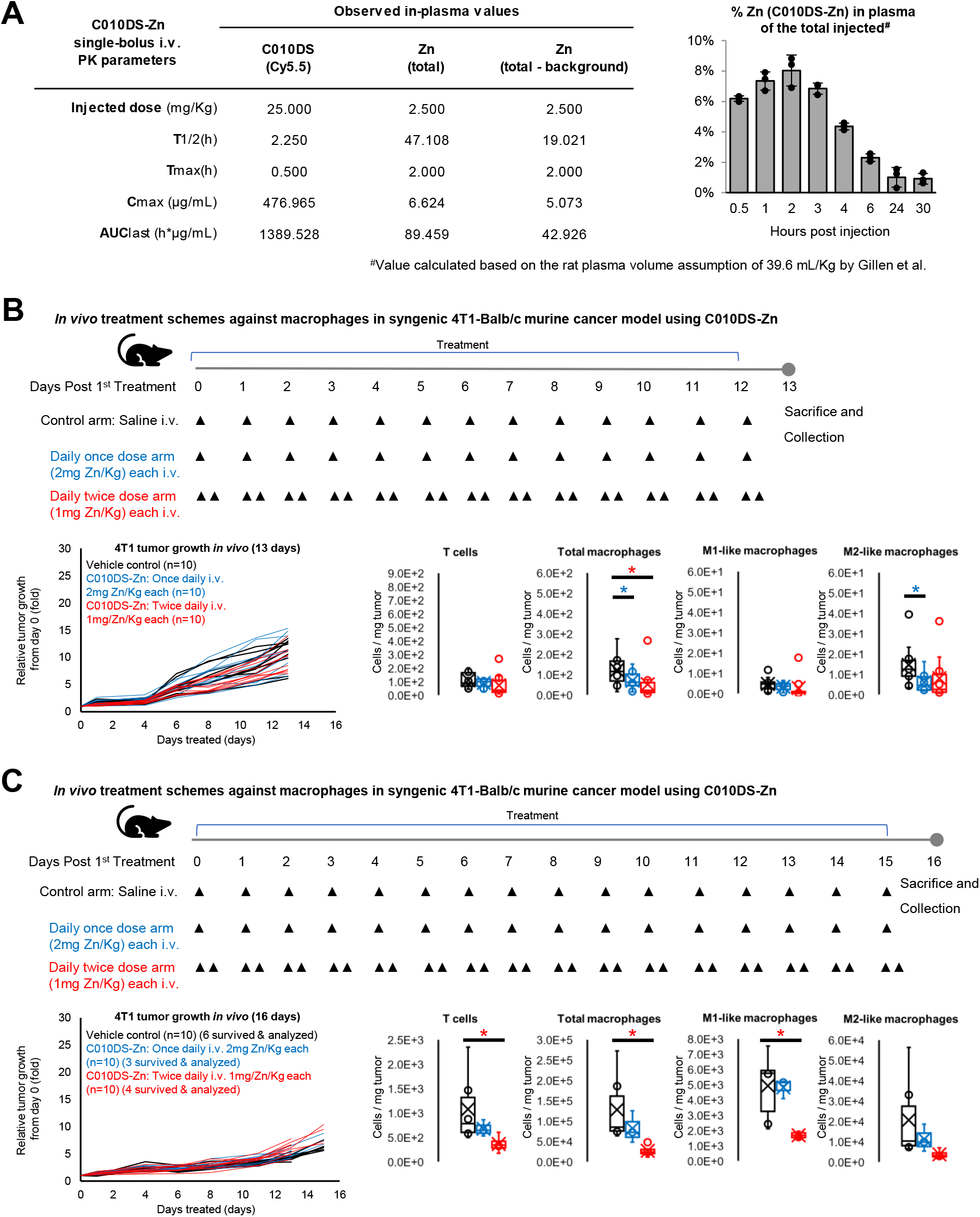
Rat plasma pharmacokinetic profiling of an intravenously dosed C010DS-Zn bolus and the *in vivo* effect of “sub-anticancer” C010DS-Zn dosing against the macrophages in the TME. **(A)** Plasma pharmacokinetic profile of C010DS-Zn that separately traced C010DS via Cy5.5 signal and zinc levels after a single intravenous bolus injection of C010DS-Zn using Sprague-Dawley Rats. Time-resolved excess plasma zinc post the C010DS-Zn injection is also plotted in % units of the total zinc injected using C010DS-Zn. **(B)** 13 days non-anticancer dosing scheme against 4T1-Balb/c model, its tumor growth kinetics, and the macrophage immune response to the treatment in the collected tumors. **(C)** 16 days non-anticancer dosing scheme against 4T1-Balb/c model, its tumor growth kinetics, and the macrophage immune response to the treatment in the collected tumors. \**p*<0.05.

To assess whether C010DS-Zn was capable of directly reducing live macrophages in the TME, we decided to test its effects on the live TME macrophage populations at lower doses that would not produce anti-tumor activity or immune initiation effects. And as 4T1 cancer model was reported to undergo TME immunity transition from early immune-suppressive state to metastasis-promoting M2-like macrophage enriched state in the late phase (*15, 16*), we decided to characterize the TME immunity at 13 days post treatment and 16 days post treatment. Consistent with the previous reports, 4T1 tumors collected on day 13 was characterized with greatly suppressed T cells and macrophage contents of both M1-like and M2-like types. In these animals, no toxicity signs meeting the General Toxicity definitions were observed during the study (**Figure 6B** and **Materials and Methods**). Those harvested on day 16, on the contrary, contained greatly increased amounts of macrophages (1E+2 in 13 days tumor vs. 1.3E+5 in 16 days tumor), mainly due to the expansion in the M2-like macrophage population, together with greatly expanded T cell content suggestive of active inflammation (**Figure 6B** and **Figure 6C**). Many of the 16-day sacrificed animals developed 50%+ tumor surface ulceration toward the end of the study, and hence were declared moribund before day 16. When treated with the lowered C010DS-Zn doses at once daily i.v. injection of 2mg Zn/Kg or twice daily injections of 1mg Zn/Kg each, the treatments led to significantly reduced macrophage content in the TME in both 13 days and 16 days treated mice. Notably in the 16 days treated groups, once daily dose of 2mg Zn/Kg led to preferential reduction in the M2-like macrophages, while twice daily doses of 1mg Zn/Kg each led to marked reduction in both M1-like and M2-like macrophages. Both C010DS-Zn treated groups of the 16 days mice also showed markedly reduced level of T cell content in the TME, suggesting reduced inflammation (**Figure 6C**).

## Discussion

Pharmaceutical utilization of parthanatos and characterization of its immunological properties in treatment against cancer have not been previously explored due to the lack of a suitable agent. In this study, we demonstrated *in vitro* that the macromolecular zinc compound C010DS-Zn uniquely induced parthanatos against the murine 4T1 cancer model at therapeutically relevant levels (**Figure 1** and **Figure 2**), which characteristically featured massive dsDNA breaks as its outcome (**Figure 3C**). C010DS-Zn’s *ex vivo* screening tests against 53 human PDX tumor fragments further corroborated that its cytotoxicity and its mode of action were highly consistent, as evidenced by the complete cytotoxicity in 53 of the 53 PDX types tested (**Figure 3**), and by the characteristic dsDNA-break intense cell death execution that was unseen in the 10% DMSO apoptosis positive controls in 50 of the 53 PDX types tested (**Figure 4**).

*In vivo*, the highest applied C010DS-Zn dosing regimen of twice daily i.v. injections at 2mg Zn/Kg each led to robust immunity initiation of the suppressed anticancer compartments in each of the model tested (**Figure 5**). Particularly against the immunosuppressive 4T1-Balb/c model featuring globally suppressed anti-tumor immunity, the treatment led to robust expansion of diverse anti-tumor immune compartments including CD4 T cells, CD8 T cells, GB+ B cells (*17*), GB+ dendritic cells (*18*), and M1-like macrophages (**Figure 5A**) to levels similar to or greater than those seen in the immunogenic CT26-Balb/c model (**Figure 5B**). This galvanizing effects of the C010DS-Zn treatment in transforming the general 4T1 TME immunity toward anti-tumor direction could be attributed to the reduced presence of CD4 Treg cells and M2-like macrophages, although the mechanism remains unclear. Based on the results of our rat PK study showing the evidence of rapid C010DS-Zn clearance by the MPS (**Figure 6A**), together with the reduced macrophage levels in the 4T1-Balb/c TME after the “non-anticancer” dose treatments independent of the 4T1 growth stages (**Figure 6B** and **Figure 6C**), we propose a direct macrophage-killing activity by C010DS-Zn as a candidate mechanism wherein the folate sidechains of the C010DS-Zn (**Figure 1A**) may be responsible for its preferential targeting of M2-like macrophages (*19*) at low-doses while the intrinsic phagocytic activity of general macrophages toward nanoparticles would lead to non-specific anti-macrophage killing at higher doses. In consideration of the core pathological roles by aberrantly expanded macrophages in the severe cases (*20*) as well as in the post-recovery cardiovascular (*21*) and autoimmune pathologies of COVID19 (i.e. Multisystem Inflammatory Syndrome in Children [MIS-C]) (*22, 23*), future investigations on the treatment potential of C010DS-Zn against such macrophage-mediated inflammatory pathologies as those seen in macrophage activation syndrome (MAS), severe COVID19, post-COVID19 myocardial damage (*21*), and MIS-C (*23*) are warranted.

In regards to the applicability and the potential of C010DS-Zn as a combination therapy with various checkpoint inhibitors, C010DS-Zn’s consistent *ex vivo* activity versus a wide variety of low-passage human solid cancer types (**Figure 4**) and the robust initiation of diverse anti-tumor immunity compartments in the 4T1-Balb/c model offer promising signs. However, in view of its low bioavailability post intravenous injection as indicated in the rat plasma PK study (**Figure 6A**), we suspect that there will be gross changes in the *in vivo* anti-cancer activity of C010DS-Zn and subsequently in the subject TME immune responses once PK-modifying formulation is applied to C010DS-Zn for its therapeutic use. With the PK-modifying formulation development already underway, we aim to test its combination treatment effects with various immunotherapeutic agents including, but not limited to, αPD1, αCTLA4, IDO inhibitors, aryl hydrocarbon receptor (AhR) inhibitors, etc.

Previously untapped, the results of our study suggest that parthanatos can be pharmaceutically accessed in potentially fighting a diverse solid cancer types using our proprietary agent C010DS-Zn like the current day chemotherapeutic agents, but with anti-cancer immune response initiation effects. As the current study is limited in many ways for fully demonstrating the therapeutic potential of C010DS-Zn against *in vivo* and real-life cancers, we plan to fill in the gaps with future follow-ups with PK-improvements, and subsequent combination therapy studies using larger numbers, longitudinal tumor re-challenge studies, and *in vivo* screening studies versus diverse PDX solid cancers in humanized immunity animals.

## Materials and Methods

### Synthesis of C005D-Zn

Synthesis of C005D (zinc salt of 32 kDa Mw γPGA conjugated to 3 folate-PEG4-NH2, 3 cRGDfK-PEG4-NH2, and 1 Cy5.5-hexane-NH2) was consigned to KRISAN Biotech (Tainan, Taiwan). Briefly, folate-PEG4-NH2 was prepared by conjugating Boc-NH2-PEG4-NH2 to the tail carboxyl group of folic acid via DIC coupling in producing the target γ-isomer, followed by TFA deprotection. The target γ-isomer was further purified by using LiChroprep RPI-18 (40-63 μm) (Merck-Millipore, Darmstadt) and subsequently lyophilized. Likewise, cRGDfK-PEG4-NH2 was prepared by conjugating 2HN-PEG4-COOH to the lysine amine of cRGDfK by EDC coupling. Once the sidechains were ready, fully protonated γPGA (32 kDa by GPC but each polymer containing 349 glutamate residues by ^1^H-NMR assessment) was conjugated to folate-PEG4-NH2, cRGDfK-PEG4-NH2, and Cy5.5-hexane-NH2 at 1:3:3:1 ratio in one-pot EDC coupling reaction using DMSO as the solvent, where 100% of the sidechains were successfully conjugated to the γPGA. The resulting peptide was then purified by solvent exchange to water using tangential flow filtration (TFF) and lyophilized to form C005D. ^1^H-NMR assessment of the C005D in DMSO-d_6_ (50g batch, #0395-069-02) sidechain contents indicated an average γPGA polymer to contain the folate-PEG4-NH2, cRGDfK-PEG4-NH2,, and Cy5.5-hexane-NH2 sidechains at the ratio of 1: 3.106: 2.607: 4.028. Finally, C005D-Zn for the biological testing of this study was prepared by mixing zinc sulfate and C005D in 30 mM HEPES-in-water of pH 7 at w/w ratio of 1:10 between the zinc ion and the γPGA backbone of the C005D.

### Synthesis of C010DS-Zn

Synthesis of C010DS (zinc salt of 32 kDa Mw γPGA conjugated to 2 folate-PEG4-NH2, 2 cRGDfK-PEG4-NH2, 10 pyrithione-S-S-PEG2-NH2, and 1 Cy5.5-hexane-NH2) was consigned to KRISAN Biotech (Tainan, Taiwan). Briefly, all sidechain preparation process and the γPGA coupling are shared with the C005D synthesis procedure above except for the pyrthione-S-S-PEG3-NH2 synthesis. Briefly, pyrthione-S-S-PEG3-NH-Boc was first synthesized by reacting dipyrithione with Boc-N-amido-PEG3-thiol in acetic acid/methanol, which was then deprotected using TFA in yielding the pyrthione-S-S-PEG3-NH2. Once the sidechains were ready, fully protonated γPGA (32 kDa by GPC but each polymer containing 349 glutamate residues by ^1^H-NMR assessment) was conjugated to folate-PEG4-NH2, cRGDfK-PEG4-NH2, pyrthione-S-S-PEG3-NH2, and Cy5.5-hexane-NH2 at 1:2:2::10:1 target ratio in one-pot EDC coupling reaction using DMSO as the solvent, where 100% of the sidechains were successfully conjugated to the γPGA. The resulting peptide was then purified by solvent exchange to water using tangential flow filtration (TFF) and lyophilized to form C010DS. ^1^H-NMR assessment of the C010DS DMSO-d_6_ (25g batch, #0523-031-33) sidechain contents indicated an average γPGA polymer to contain the folate-PEG4-NH2, cRGDfK-PEG4-NH2, pyrthione-S-S-PEG3-NH2, and Cy5.5-hexane-NH2 sidechains at the ratio of 1: 1.53: 1.62: 9.04: 1.18. Finally, C010DS-Zn for the biological testing of this study was prepared by mixing zinc sulfate and C010DS in 30 mM HEPES-in-water of pH 7 at w/w ratio of 1:10 between the zinc ion and the γPGA backbone of the C010DS.

### Cell preparations *in vitro* tests (Consigned to Invitrocue)

Mouse mammary carcinoma 4T1 cells (ATCC^®^ CRL-2539^™^) and mouse colorectal carcinoma CT26 cells (ATCC^®^ CRL-2638^™^) were cultured in RPMI Medium 1640 (61870-036, Gibco) supplemented with 10% Foetal Bovine Serum (FBS; HyClone, SH3007103) and 1% Penicillin-Streptomycin (P/S; Sigma Aldrich, P4333). Cells were incubated at 37°C with 5% CO_2_. Cells were passaged upon 80-90% confluency using 0.25% trypsin-EDTA. Manual cell count was performed using haemocytometer.

### Drug preparation for IC50 determination using LDH assay

Zinc sulfate heptahydrate (Sigma Aldrich, Z0251), was weighed and dissolved in sterilized water to prepare 10 mg/mL of stock solutions. C005D-Zn and C010DS-Zn stocks were solubilized in 30 mM pH 7 HEPES buffer at concentrations of 1mg Zn/mL. PJ34 (Santacruz Biotech Cat# sc-204161A) stock solution was prepared at 2mM concentration in water.

### LDH-Glo^™^ Cytotoxicity Assay (Consigned to Invitrocue)

Lactate dehydrogenase (LDH) assay was performed by the kit manufacturer’s LDH-Glo^™^ Cytotoxicity Assay protocol (Promega). Briefly, cells were seeded into 96-well plates at a seeding density of 7,500 cells per well and incubated overnight at 37°C with 5% CO_2_. The cells were then treated with test compound or vehicle control for 24 h prior to the assay. Media post-treatment was collected and diluted 50 times with LDH storage buffer before the measurement of LDH released. LDH detection reagent was then added to the diluted media at 1:1 ratio and luminescence reading was measured using Infinite^®^ 200 microplate reader (Tecan).

### Confocal Imaging studies (Consigned to Invitrocue)

The terminal deoxynucleotidyl transferase dUTP nick end labeling (TUNEL), Apoptosis-inducing factor (AIF) and Hoechst 33342 triple staining was used to detect the DNA fragmentation and AIF translocated to nucleus. 4T1 cells were plated at 20,000 cells per well in 96 well black imaging plates and incubated overnight at 37°C with 5% CO_2_. Cells were then treated with test compounds at concentrations shown in Table 1 for 3 hours, in presence or absence of 0.5 μM PJ34. Cells were fixed with 4% paraformaldehyde and permeabilized with 0.25% of Triton X-100 before staining with Click-iT^®^ TUNEL Alexa Fluor^®^ Imaging Assay (Invitrogen, C10245), Hoechst 33342 (Invitrogen, C10247, dilution factor 1:2000) and AIF (Clone E-1) antibody Alexa Fluor^®^ 594 (Santa Cruz, sc-13116, dilution factor 1:400) at 4°C overnight. Images were captured with a confocal laser scanning microscope (LSM 700, Zeiss, US) at 405nm, 488 nm and 594nm, respectively. The AIF and TUNEL translocation to nucleus were further analyzed using the CircAvgInten function of the CellInsight^™^ CX7 LZR High-Content Screening (HCS) Platform (ThermoFisher, US).

### Poly ADP-Ribose (PAR) ELISA Assay (Consigned to Invitrocue)

Poly ADP-ribose (PAR) level in cells was evaluated using the PAR ELISA Kit (Cell Biolabs, XDN-5114). Briefly, 4T1 cells were seeded at a density of 30,000 cells per well on a 24-well plate and incubated overnight at 37°C with 5% CO_2_. Cells were then treated with 2.5, 5, 10, 20 μM of C010DS-Zn in presence or absence of 0.5 μM PJ34 inhibitor. After 24h of treatment, cells were lysed using RIPA lysis buffer (Sigma-Aldrich, R0278) and supernatant was collected after centrifugation at 10,000 g for 10min at 4°C. PAR ELIZA assay was performed according to manufacturer’s kit instructions. Absorbance was measured at 450 nm using Infinite^®^ 200 microplate reader (Tecan, Switzerland).

### *In vitro* characterization of A5 and PI uptake using flow cytometer (Consigned to Invitrocue)

Cell death mechanism was analyzed using Annexin V-FITC apoptosis detection kit protocol (eBioscience, BMS500FI/300) according to the manufacturer’s instructions. Briefly, 4T1 cells were seeded at a density of 50,000 cells per well on a 24-well plate and incubated overnight at 37°C with 5% CO_2_. Cells were then were treated with test compounds or vehicle control at concentrations shown in Table 2. Cells were harvested, washed, and stained with Annexin V and propidium iodide (PI), and analyzed on a CytoFLEX LX Flow Cytometer (Beckman Coulter Life Sciences, US) with a 488 nm argon ion laser and a 635 nm red diode laser. Flow cytometric data acquisition and analysis were conducted using FlowJo v10.7.1 (BD Biosciences, US).

### TME immune cell content characterization in the tumors harvested from the *in vivo* anti-tumor treatment test against 4T1-Balb/c and CT26-Balb/c tumor models (Consigned to Invivocue)

#### Mice and animal care

Female BALB/cAnNTac (BALB/c) were purchased from InVivos Singapore. All mice were bred and kept under pathogen-free conditions in Biological Resource Centre, Agency for Science, Technology and Research, Singapore (A*STAR) on controlled 12 hour light-dark cycle. All experiments and procedures were approved by the Institutional Animal Care and Use Committee (IACUC# 191485) of A*STAR in accordance with the guidelines of Agri-Food and Veterinary Authority and the National Advisory Committee for Laboratory Animal Research of Singapore.

#### CT26 and 4T1 syngeneic mouse model

CT26 colon carcinoma cells and 4T1 mammary carcinoma were purchased from ATCC and cultured in RPMI 1640 (Gibco) with 10% (vol/vol) FBS. 1 × 10^5^ of CT26 cells (50μl) or 3 × 10^4^ 4T1 cells (50μl) was subcutaneously injected into shaved right flank of each BALB/c mouse (8-10 wko). Drug dosing was initiated when tumor volume reaches approximately 50-100 mm3. CT26 tumor bearing mice were administered with either C010DS-Zn (2mg/kg, daily intravenously), C010DS-Zn (2mg/kg, daily twice intravenously) or saline/vehicle control (5μl per g of body weight, daily intravenously). The CT26 tumor bearing mice were administered with drugs for 12 days followed by two days of observation before sacrificed for further analysis. Similarly, 4T1 tumor bearing mice were administered with either C010DS-Zn (2mg/kg, daily intravenously), C010DS-Zn (1mg/kg, daily twice intravenously) or saline/vehicle control (5μl per g of body weight, daily intravenously). The 4T1 tumor bearing mice were administered with drugs for 10 days before sacrificed for further analysis.

#### Determination of tumor size

The length and width of tumor was measured using caliper and the tumor volume was calculated using the formula: Tumor volume = (length × width^2^) × 0.5.

#### Tumor infiltrating immune cells isolation

Tumor-infiltrating cells were isolated and enriched using mouse tumor dissociating kits (Militenyi), following manufacturer’s instructions. The isolated tumor infiltrating immunce cells were centrifuged in 35% v/v Percoll solution (GE Healthcare) and RPMI (Sigma-Aldrich) prior to staining procedures.

#### Immunophenotyping of TME leukocytes

Live immune cells from tumor were determined by staining with live/dead fixable blue dead cell stain kit (Life Technologies) for 30 min prior to cell-specific marker staining, following manufacturer’s protocol. Staining panel for CT26 and 4T1 mouse syngeneic model is as follows: anti-mouse CD4 (BD Biosciences, cat#563790, clone# GK1.5), anti-mouse CD44 (BD Biosciences, cat#612799, clone#IM7), anti-mouse PD-1 (Biolegend, cat#109121, clone#RMP1-30), anti-mouse CD19 (Biolegend, cat# 115546, clone#6D5; BD Biosciences, cat#563333, clone#1D3), anti-mouse CD45 (BD Biosciences, cat#563053, clone#30-F11), anti-mouse CD62L (BD Biosciences, cat#564108, clone#MEL-14), anti-mouse Ly6C/Ly6G (BD Biosciences, cat#740658, clone# RB6-8C5), anti-mouse CD335 (BD Biosciences, cat#741029, clone#29A1.4), anti-mouse Granzyme B (Biolegend, cat#396404, clone#QA18A28), anti-mouse FoxP3 (BD Biosciences, cat#560408, clone#MF23), anti-mouse CD8 (BD Biosciences, cat#552877, clone#53-6.7), anti-mouse CD3 (Biolegend, cat#100236, clone#17A2; BD Biosciences, cat#564010, clone#17A2), anti-mouse CD25 (Biolegend, cat#101918, clone#3C7), anti-mouse CD24 (BD Biosciences, cat#744471, clone#M1/69), anti-mouse I-A/I-E (BD Biosciences, cat#748708, clone#2G9), anti-mouse F4/80 (BD Biosciences, cat#565411, clone#T45-2342), anti-mouse Ly6C (Biolegend, cat#128033, clone#HK1.4), anti-mouse CD206 (Biolegend, cat#141723, clone#C068C2), anti-mouse CD11c (Biolegend, cat#117308, clone#N418), anti-mouse Ly6G (BD Biosciences, cat#560601, clone#1A8), anti-mouse CD11b (BD Biosciences, cat#563312, clone#M1/70) and anti-mouse CD115 (Biolegend, cat#135532, clone#AFS98). Cells were first stained with surface markers for 30 min at room temperature. After which, cells were washed in FACs buffer containing phosphate buffered saline (PBS), 0.2% bovine serum albumin (GE Healthcare Life Sciences) and 0.05% sodium azide (Merck), fixed and permeabilised (Mouse Foxp3 Buffer Set from BD Biosciences) before stained with intracellular markers for 30 min at room temperature. Cells were then washed and resuspended in FACs buffer for flow cytometry data acquisition. Data was acquired using a LSR II flow cytometer (BD Biosciences) with FACSDiva software, and analyzed using FlowJo software (version 10; Tree Star Inc). Gating strategy employed at Invivocue for the TME leukocyte characterization is shown in **Figure S1** (T cell and B cell panel) and **Figure S2** (myeloid cell panel).

### *In vivo* anti-macrophage treatment test against 4T1-Balb/c tumor models (Consigned to Champions Oncology)

#### Mice and animal care

Female BALB/c were purchsed from Taconic and acclimated for 3 days. All mice were bred and kept under pathogen-free conditions in Champions Oncology facility on controlled 10-14 hour light-dark cycle at 20-23 °C and 30-70% humidity. Only the animals meeting the minimum 6-8 weeks of age and 18 grams of weight at the time of dosing were used for the study. All experimental procedures were performed according to the guidelines of the Institutional Animal Care and Use Committee (IACUC) of Champions Oncology.

#### 4T1-Balb/c mouse study design

Pre-study animals (female, BALB/c) were injected with 0.1 mL of 300,000 4T1 cells subcutaneously, in the left flank. Pre-study tumor volumes were recorded for each experiment beginning four to five days after injection. Tumor growth was monitored beginning 4-5 days after implantation using digital calipers and the tumor volume (TV) was calculated using the formula (0.52 × [length × width^2^]). When tumors reach an average tumor volume of 50-150 mm^3^ animals were matched by tumor volume into treatment or control groups (n=10/group) to be used for dosing and were treated intravenously with vehicle QD×12 or QD×15, Xylonix Agent, C010DS-Zn at 1 mg Zn/kg BID×12 or BID×15 and Xylonix Agent, C010DS-Zn at 2 mg Zn/kg QD×12 or QD×15. After the initiation of dosing on Day 0, animals were weighed three times per week using a digital scale and TV was measured three times per week and on the final day of study. A final weight was taken on the day the study reached end point or if animal was found moribund, if possible. Animals exhibiting >10% weight loss when compared to Day 0 was provided DietGel^®^ ad libitum. Any animal exhibiting >20% net weight loss for a period lasting 7 days or if mice display >30% net weight loss when compared to Day 0 was considered moribund and euthanized. The study endpoint was when the vehicle control mean tumor volume reached 1500 mm3. If this occured before Study Day 13 (i.e.14 days on study) or Day 16 (i.e.17 days on study), treatment groups and individual mice may be dosed and measured up to Study Day 13 or Study Day 16. Tumors can start developing ulceration at early time points; triple antibiotic ointment would be applied three times per week on all tumors after tumor volume measurements. Mice with ulceration greater than 50% of the tumor total surface area or an impact on overall health and well-being will be euthanized.

#### General toxicity

Beginning on Day 0, animals were observed daily and weighed thrice weekly using a digital scale; data including individual and mean gram weights, mean percent weight change versus Day 0 (%vD0) were recorded for each group and %vD0 plotted at study completion. Animals exhibiting ≥ 10% weight loss when compared to Day 0, if any, were provided with DietGel^™^ (ClearH2O^®^, Westbrook, ME) ad libitum. Animal deaths, if any, were recorded. groups reporting a mean loss of %vD0 >20 and/or >10% mortality were considered above the maximum tolerated dose (MTD) for that treatment on the evaluated regimen. Additional study toxicity endpoints were mice found moribund or displayed >20% net weight loss for a period lasting 7 days or if the mice displayed >30% net weight loss. Maximum mean %vD0 (weight nadir) for each treatment group was reported at study completion.

#### Determination of tumor size

The length and width of tumor was measured using caliper and the tumor volume was calculated using the formula: Tumor volume = (length × width^2^) × 0.5.

#### Immunophenotyping of TME leukocytes

Harvested 4T1 tumors from the Balb/c mice were weighed and placed into MACS media, then stored on wet ice until processed for flow cytometry. The tumors were then dissociated using MACS Miltenyi Biotech Tumor Dissociation Kit (Cat#130-096-730) per manufacturer protocol, and the dissociated tumors were stained by Champion Oncology’s standard staining protocol for surface and intracellular markers that consisted of mCD25-BV421 (PC61), mF4/80-BV510 (BM8), mCD45-BV605 (30-F11), mCD11b-BV711 (M1/70), mCD44-BV786 (IM7), mCD8a-AF488 (53-6.7), mMHC0II-PerCP Cy5.5 (M5/114.15.2), mFoxP3-PE (FJK-16s), mCD3-PE Cy7 (17A2), mCD206-AF647 (C068C2), mCD4-APC Fire 750 (RM4-5), and FVS700 (viability). Gating strategy employed at Champions Oncology for the TME immune cell characterization is shown **Figure S3**.

### Imaging-based *Ex vivo* screening of C010DS-Zn cytotoxicity against low passage 53 human solid cancer PDX fragments (Consigned to Champions Oncology)

Champions Oncology (Hackensack, NJ) operates a bank of low passage human cancer patient tumor specimens that have been catalogued with the patient treatment information as well as the NGS-sequenced genetic information (Champions TumorGraft Models^™^) that were available for *ex vivo* cytotoxicity screening studies on consignment basis. Initially 75 human solid cancer types were selected from triple negative breast cancer, non-triple negative breast cancer, colorectal cancer, non-small cell lung cancer, prostate cancer, pancreatic cancer, and sarcoma with even distribution of TMB and/or MSI scores. Due to quality control issues, 53 of the 75 selected models were completed as originally scheduled and hence included in the current study. Briefly for each model, cryopreserved tumor samples (200-300 mg or more depending on the need) was dissociated manually using Gentle MACS Dissociator and human Tumor Dissociation Kit (Miltenyi, Bergisch Gladbach). After the preparation, tumor fragments were cultured overnight, or in certain cases, for up to 7 days before filtering out large fragments using a 500 μm filter followed by a 200 μm filter. Flow-through was cultured in Cell-Tak coated low volume flat bottom 384-well assay plates. Using 50 μg Zn/mL as the top C010DS-Zn concentration prepared from the Xylonix-provided C010DS-Zn stock at 1mg Zn/mL, serial half-log dilution scheme was used in testing the cytotoxicity response curve against each model with 4 replicates (n=4), a negative control group using 0.1% DMSO, and a positive control group using 10% DMSO. To avoid edge-effect artifacts in each plate, PBS was plated in the perimeter wells. The plated tumor fragments were then treated for 120 hours (5 days), at which point their cell viability was read using CellTiter-Glo assay (Promega, Madison). Additionally, fluorescence imaging data of the treated tumor fragments were obtained using Hoeschst, Cell Tracker Green, and pγH2A.X for the 2 highest concentrations of the tested agents, as well as for the negative and the positive controls. Viability checks were performed using CellTiter-Glo assay upon plating, and at the assay endpoint. Processed C010DS-Zn IC50 value versus each model, as well as the representative imaging data were reported to back to Xylonix.

### In-rat PK study (Consigned to Pharmaron)

#### Rats and animal care

Male naïve Sprague-Dawley rats [Crl:CD(SD)IGS] (SPF/VAF) were purchased from Vital River Laboratory Animal Technology (Beijing) and acclimated for at least 5 days. All animals were bred and kept in cages (2-3 animals per cage) under pathogen-free conditions in Pharmaron as managed by the office of Laboratory Animal Medicine Department of Pharmaron. Controlled 12-12 hour light-dark cycle at 20-26 °C and 40-70% relative humidity under 15 air changes per hour per room were applied. Only the animals meeting the age between 6-9 weeks and 200-220g of weight at the time of first dosing were used for the study. All experimental procedures were reviewed, approved and performed according to the guidelines of the Pharmaron Institutional Animal Care and Use (IACUC). Pharmaron has been certified by the International Experimental Animal Assessment Committee (AAALAC).

#### PK study design

27 male Sprague-Dawley rats were randomly selected from total 31 animals by body weights and assigned to the study using a computerized procedure using Provantis program to achieve body weight balance with respect to group assignment with no significant weight difference among 9 groups for each time point (n=3). For the PK study, a single intravenous bolus injection of pH 7 C010DS-Zn in 30 mM HEPES-water solution was used as the dosing method at the dose volume of 5 mL/kg targeting the C010DS-Zn dose level containing 2.5 mg Zn/Kg or 25 mg γPGA-H/Kg. The absolute dose volume for each animal was calculated using Provantis 9.3.1 software based on the most recently measured body weight. After the C010DS-Zn single bolus intravenous injection, blood samples were collected using K_2_EDTA tube for each time-point group of 3 animals (n=3) via puncture of abdominal aorta, 0.5 mL for LC-MS/MS (metabolite analysis), 0.5 mL for ICP-MS (zinc analysis), and 1.0 mL for fluorescence (Cy5.5 analysis). The tubes containing the whole blood were then inverted several times and placed on wet ice/ice bag, and centrifuged at 2,000g for 10 mins at 4 °C to obtain cell-free plasma. For LC-MS/MS samples, the plasma samples were further treated with “methanol:5 mM ammonium acetate solution” (50:50, v/v) containing 2 mg/mL of 2-mercaptoethanol (50:5) and 2% TFA after collection for stabilization. The stabilized LC-MS/MS samples and ICP-MS were then shipped to the clinical analytical testing laboratory at Pharmaron (BDA) for analysis. The fluoresence analysis samples were analyzed immediately at the Immunology lab of Pharmaron-TSP.

#### Rat serum PK data analysis

The cell-free serum PK parameters were calculated using WinNonlin (Phoenix^™^ version 8.1 or higher).

### Statistics

All statistical comparisons between any 2 groups in this study was performed by using the T.Test function of Excel (Microsoft^®^, Seattle) assuming single-tailed comparison between two groups of equal variances.

## Author contribution

Invention of proprietary substances used, study funding, and study contractor sourcing – JFC

Study design and insights – JFC, ZH, WMK, and QF

Lab works – ZH and WMK

Data analysis – JFC, ZH, and WMK

Manuscript writing – JFC

## Acknowledgements

We thank Ms. Susan Chia at Invitrocue for technical assistance on the *in vitro* works performed for this study. We thank the liaison officer and study coordinator Ms. Veena Jagannathan and Mr. Christopher Noakes at Champions Oncology for the design and selection of the human PDX cancer models, and for the *in vitro*, *in vivo*, and *ex vivo* works performed for this study. We thank the coordination officers Dr. Tina Yen and Mr. Jason Yang, synthesis chemist Mr. Elvis Yu and Mr. Longhu Wang, and R&D director Dr. Bret Wu at KRISAN biotech for the manufacturing synthesis development works on C005D-Zn and C010DS-Zn. We thank Dr. Xiaonan Liang of Pharmaron for the PK study, and Dr. Robert Jiang for numerous inputs on toxicology and PK study designs and interpretations. Special thanks to the CEO Dr. Yu Pei Wang of KRISAN Biotech and the CEO Dr. Steven Fang of Invitrocue for special administrative arrangements that enabled this study.

