## Supplemental figures for "Parthanatos-inducing zinc agent C010DS-Zn elicits anti-tumor immune responses involving T cells and macrophages *in vivo*"

Panel 1 –
T and B cell gating strategy used at Invivocue™ for the anti-tumor treatment experiments against 4T1-Balb/c and CT26-Balb/c syngenic models

(Figure S1)

| Panel 1: Xylonix 4T1 Syngenic Study - NK and T panel | | | | |
|---|---|---|---|---|
| No. | Fluorochrome | Antigen | Cat# | Description |
| 1 | BUV395 | CD4 | BD 563790 | T helper |
| 2 | DAPI | L/D | LifeTech #L34962 | Live/Dead |
| 3 | BUV737 | CD44 | BD 612799 | T memory |
| 4 | BV421 | PD-1/CD279 | Biolegend 109121 | T exhaustion marker |
| 5 | BV510 | CD19 | BD 562956 | B cells |
| 6 | BV605 | mCD45 | BD 563053 | pan immune cell marker |
| 7 | BV650 | CD62L | BD 564108 | T memory |
| 8 | BV711 | Ly6C/Ly6G | BD 740658 | myeloid cells exclusion |
| 9 | BV786 | CD335 | BD 741029 | NK cells |
| 10 | FITC/AF488 | Granzyme B (Intracellular) | Biolegend 396404 | anti-tumor cytotoxicity |
| 11 | PE | Foxp3 (Intracellular) | BD 560408 | Treg |
| 12 | PeCy7 | CD8 | BD 552877 | T cytotoxic |
| 13 | APC/AF647 | CD3 | Biolegend 100236 | pan T cell marker |
| 14 | APC-H7 | CD25 | Biolegend 101918 | Treg |

### Schematic Diagram of Gating Strategy

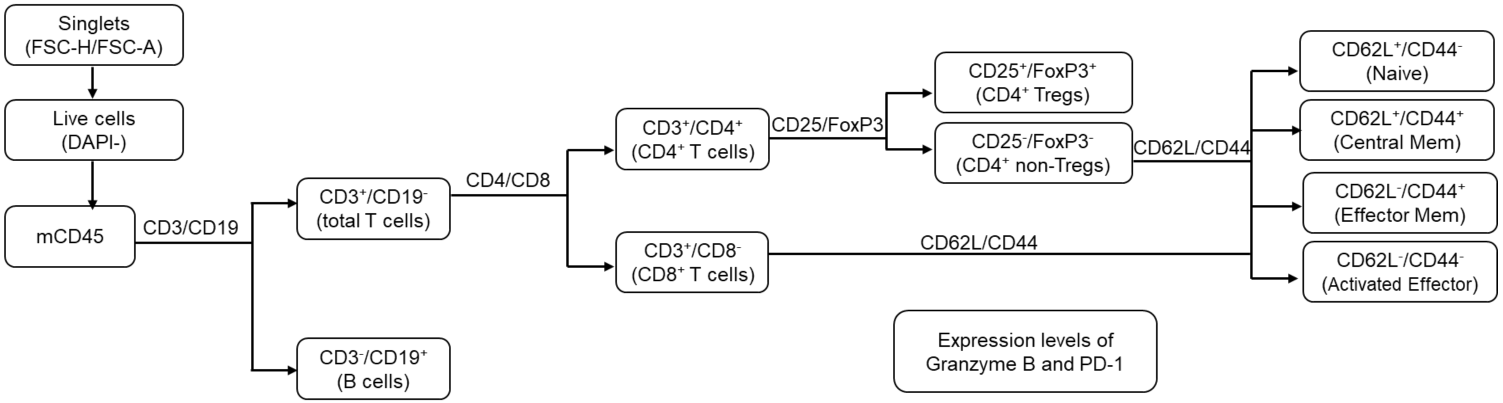


*Expression levels of Granzyme B and PD1 were determined in key subsets.

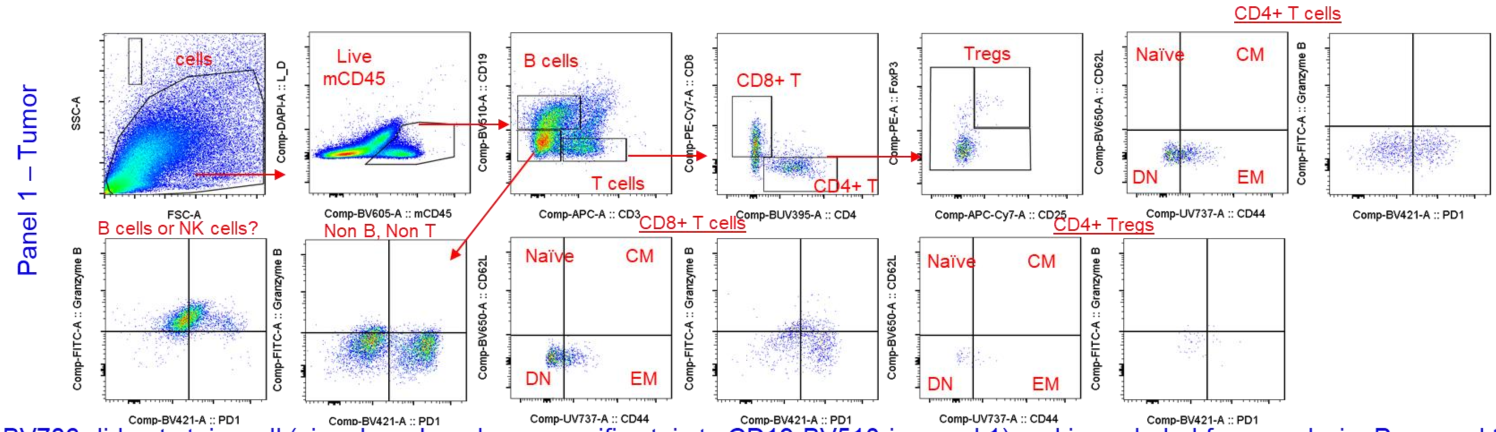


CD335-BV786 did not stain well (signal weak and non-specific stain to CD19-BV510 in panel 1) and is excluded from analysis. Proposed to change CD335 to NKG2ACE for future staining if NK cells is of interest

Panel 2 –
Myeloid cell gating strategy used at Invivocue™ for the anti-tumor treatment experiments against 4T1-Balb/c and CT26-Balb/c syngenic models

(Figure S2)

| Panel 2: Xylonix - Myeloid panel | | | | |
|---|---|---|---|---|
| No. | Fluorochrome | Antigen | Cat# | Description |
| 1 | BUV395 | CD24 | BD 744471 | Mac/ DC cell delination |
| 2 | DAPI | L/D | LifeTech #L34962 | Live/Dead |
| 3 | BUV737 | MHCII | BD 748708 | M1 mac/DC |
| 4 | BV421 | F4/80 | BD 565411 | pan macrophage |
| 5 | BV510 | Ly6C | Biolegend 128033 | mono/neutrophils/m-MDSC |
| 6 | BV605 | mCD45 | BD 563053 | pan immune cell marker |
| 7 | BV650 | CD206 (Intracellular) | Biolegend 141723 | M2 mac |
| 8 | BV786 | CD3 | BD 564010 | T, B, NK cell exclusion |
| 9 | BV786 | CD335 | BD 741029 | T, B, NK cell exclusion |
| 10 | BV786 | CD19 | BD 563333 | T, B, NK cell exclusion |
| 11 | FITC/AF488 | Granzyme B (Intracellular) | Biolegend 396404 | anti-tumor cytotoxicity |
| 12 | PE | CD11c | Biolegend 117308 | pan T cell marker |
| 13 | PeCy7 | Ly6G | BD 560601 | DC |
| 14 | APC/AF647 | CD11b | BD 553312 | pan myeloid lineage |
| 15 | APC-H7 | CD115 | Biolegend 135532 | MDSC-associated |

### Schematic Diagram of Gating Strategy

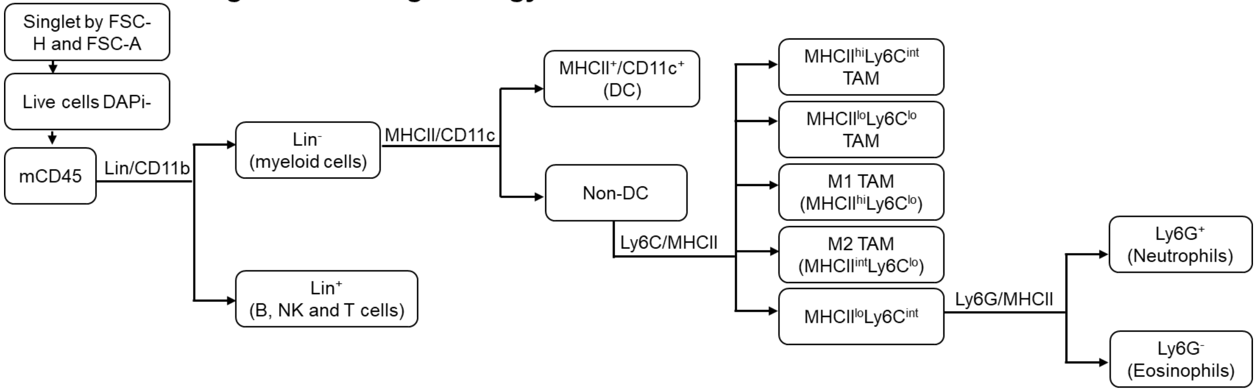


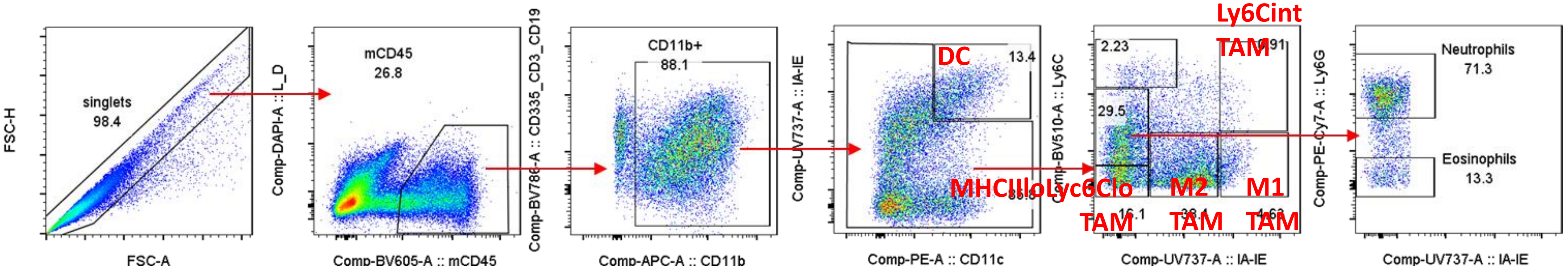

Panel 2 – Myeloid cell gating strategy used at Invivocue™ for the anti-tumor treatment experiments against 4T1-Balb/c and CT26-Balb/c syngenic models

(Figure S3)

### Antibody selection panel

| Channel | Antibody | Conjugate | Clone |
|---|---|---|---|
| V1 | mCD25 | BV421 | PC61 |
| V2 | mF4/80 | BV510 | BM8 |
| V4 | mCD45 | BV605 | 30-F11 |
| V6 | mCD11b | BV711 | M1/70 |
| V7 | mCD44 | BV786 | IM7 |
| B1 | mCD8a | AF488 | 53-6.7 |
| B3 | mMHC-II | PerCP Cy5.5 | M5/114.15.2 |
| YG1 | mFoxP3 | PE | FJK-16s |
| YG4 | mCD3 | PE Cy7 | 17A2 |
| R1 | mCD206 | AF647 | C068C2 |
| R2 | Viability | FVS700 | N/A |
| R3 | mCD4 | APC Fire 750 | RM4-5 |

### Gating strategy summary

| Description | Parent Gate | Gate | Name | Units |
|---|---|---|---|---|
| Scatter | All events | FSC-A v SSC-A | Cells | |
| FSC singlets | Cells | FSC-H v FSC-A | FSC singlets | |
| SSC singlets | FSC singlets | SSC-H v SSC-A | SSC singlets | |
| Live | SSC singlets | SSC-A v FVS700 | Live | |
| CD45 | Live | SSC-A v CD45+ | CD45 | %, Events, cells/mg of tumor |
| Total T cells | CD45 | CD3+ v SSC-A | Total T cells | %, Events, cells/mg of tumor |
| CD4 T cells | Total T cells | CD4+ v CD8- | CD4 T cells | %, Events, cells/mg of tumor |
| CD4+CD25+ T cells | Total T cells | CD4+ v CD25+ | CD4+ CD25+ T cells | %, Events, cells/mg of tumor |
| CD4 T reg | CD4 T cells | CD25+ v FoxP3+ | CD4 Tregs | %, Events, cells/mg of tumor |
| CD4+ CD44+ T cells | CD4 T cells | CD4+ v CD44+ | CD4+ CD44+ T cells | %, Events, cells/mg of tumor |
| CD8 T | Total T cells | CD4- v CD8+ | CD8 T cells | %, Events, cells/mg of tumor |
| CD8+ CD25+ T cells | CD8 T cells | CD8+ v CD25+ | CD8+ CD25+ T cells | %, Events, cells/mg of tumor |
| CD8+ Treg | CD8 T cells | CD25+ v FoxP3+ | CD8+ Treg | %, Events, cells/mg of tumor |
| CD8+ CD44+ cells | CD8 T cells | CD8+ v CD44+ | CD8+ CD44+ cells | %, Events, cells/mg of tumor |
| Macrophage | CD45+ | F4/80+CD11b+ | Macs | %, Events, cells/mg of tumor |
| M1 Macs | Macs | MHCII+CD206- | M1 Mac | %, Events, cells/mg of tumor |
| M2 Macs | Macs | MHCII-CD206+ | M2 Mac | %, Events, cells/mg of tumor |
| CD11b+ | CD45+ | SSC-a v CD11b | CD11b+ | %, Events, cells/mg of tumor |

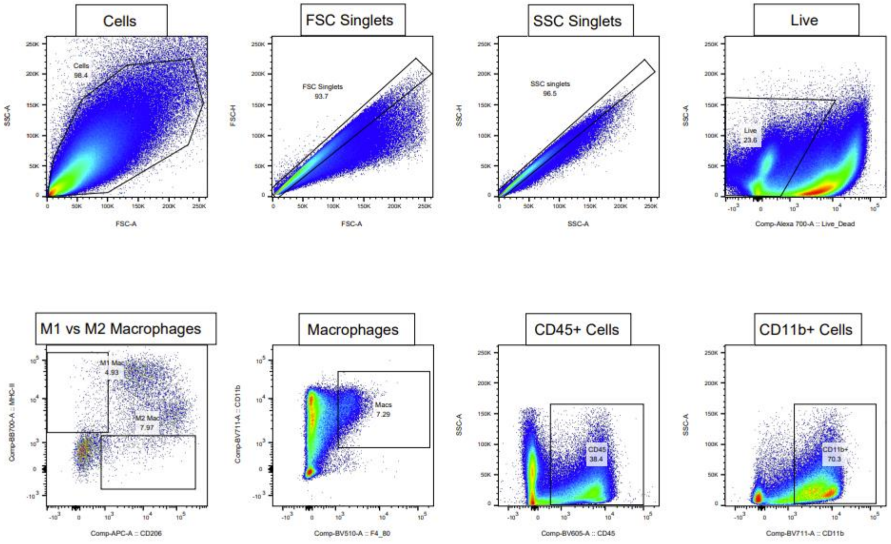


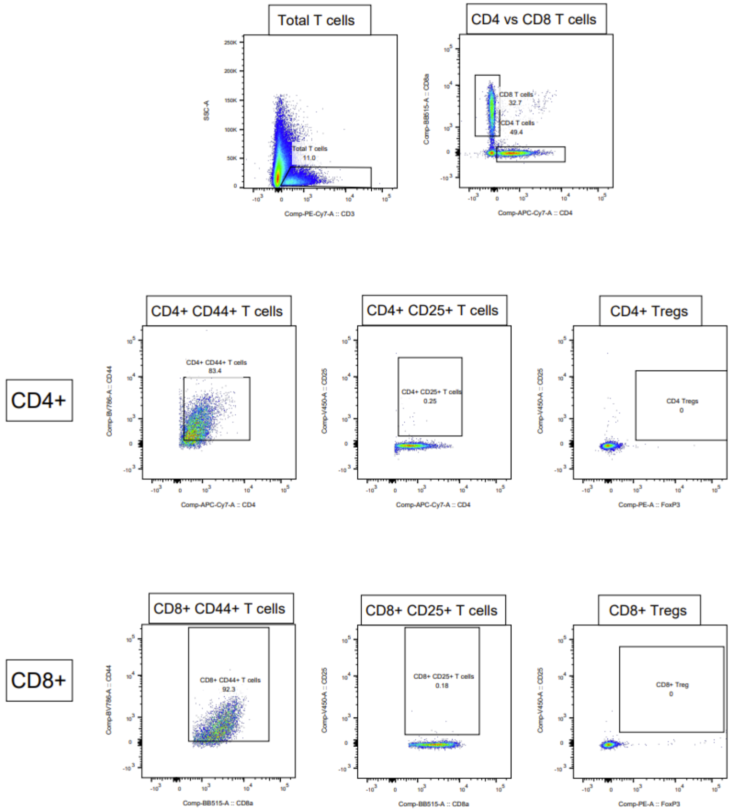
